# Plasticity in stomatal behavior across a gradient of water supply is consistent among field-grown maize inbred lines with varying stomatal patterning

**DOI:** 10.1101/2021.10.28.466255

**Authors:** Risheng Ding, Jiayang Xie, Dustin Mayfield-Jones, Yanqun Zhang, Shaozhong Kang, Andrew D.B. Leakey

## Abstract

Stomata regulate leaf CO_2_ assimilation (*A*) and water loss. The Ball–Berry and Medlyn models predict stomatal conductance (*g*_s_) with a slope parameter (*m* or *g*_1_) that reflects sensitivity of *g*_*s*_ to *A*, atmospheric CO_2_ and humidity, and is inversely related to water use efficiency (WUE). This study addressed knowledge gaps about what the values of *m* and *g*_1_ are in C_4_ crops under field conditions, as well as how they vary among genotypes and with drought stress. *m* and *g*_1_ were unexpectedly consistent in four inbred maize genotypes across a gradient of water supply. This was despite genotypic variation in stomatal patterning, *A* and *g*_s_. *m* and *g*_1_ were strongly correlated with soil water content, moderately correlated with pre-dawn leaf water potential (Ψ_pd_), and weakly correlated with midday leaf water potential (Ψ_md_). This implied that *m* and *g*_1_ respond to long-term water supply more than short-term drought stress. The conserved nature of *m* and *g*_1_ across anatomically diverse genotypes and water supplies suggests there is flexibility in structure-function relationships underpinning WUE. This evidence can guide simulation of maize *g*_s_ across a range of water supply in the primary maize growing region and inform efforts to improve WUE.

**Summary statement:** Parameter values for models simulating stomatal conductance were unexpectedly consistent for anatomically and physiologically diverse genotypes of the model C_4_ crop maize when they were grown across a range of water supplies in the field.

## Introduction

Stomata on leaf surfaces regulate the exchange and trade-off of carbon and water between plants and the atmosphere while responding to environmental and physiological signals (Berry, Beerling, & Franks, 2010; Hetherington & Woodward, 2003). Stomatal conductance (*g*_s_) is a key determinant of leaf, plant, canopy, ecosystem, regional and global fluxes of water, carbon, and energy (Bonan, Williams, Fisher, & Oleson, 2014; Franks, Berry, Lombardozzi, & Bonan, 2017). Therefore, stomatal conductance is a key regulator of crop performance as well as the biogeochemistry of natural and managed ecosystems (Leakey et al. 2019). And, a mathematical representation of *g*_s_ is one fundamental component of models of plant and ecosystem function (Lawrence et al., 2019; Oleson et al., 2010; Sellers et al., 1997).

Two of the most widely used models of *g*_s_ are the Ball-Berry (BB) model (Ball, Woodrow, & Berry, 1987) and Medlyn (MED) model (Medlyn et al., 2011). Both the BB and MED models describe *g*_s_ as a function of the rate of net photosynthetic carbon dioxide assimilation (*A*) and atmospheric carbon dioxide concentration at the leaf surface (*C*_s_), along with either atmospheric relative humidity (*H*_s_) or vapor pressure deficit (*D*_s_) at the leaf surface. Stomatal behavior in response to these drivers is captured in terms of a slope parameter (*m* or *g*_1_) and intercept parameter (*g*_0_ or *g*_0M_). The slope parameter *m* reflects the sensitivity of *g*_s_ to changes in *A*\**H*_s_/*C*_s_ (hereafter referred to as the BB Index), and *g*_1_ to *A*/(*C*_s_√*D*_s_) (hereafter referred to as the MED Index). Biologically, *m* and *g*_1_ are an inverse of intrinsic water use efficiency (iWUE, *A*/*g*_s_) for given set of environmental conditions, e.g., fixed *H*_s_ or *D*_s_ and *C*_s_ (Leakey, Bernacchi, Ort, & Long, 2006; Wolz, Wertin, Abordo, Wang, & Leakey, 2017). As an extension of these basic models, there have been numerous formulations to incorporate the effects of water stress on *g*_s_, all of which are based on empirical functions describing the response of *g*_*s*_ model parameters to variation in soil or plant water status (Damour, Simonneau, Cochard, & Urban, 2010).

Recent studies have highlighted the importance of understanding how the slope parameters of *g*_s_-models vary among plants and across environmental gradients (Franks *et al.*, 2018; Lin et al., 2015; Miner and Bauerle 2017; Wolz *et al.*, 2017). But, despite the importance of *g*_s_ and models of *g*_s_, knowledge gaps remain about: (1) how the slope parameters of stomatal conductance models vary within crop species; (2) whether there are significant genotype by environment interactions; and (3) how stomatal patterning on the epidermis may influence model representations of *g*_s_.

There is clear evidence for substantial variability in the parameters of stomatal conductance models within and across different plant functional types (Franks et al., 2018; Miner and Bauerle 2017; Wolz *et al.*, 2017). And, the limited available data suggest that intraspecific variation can be as great as interspecific variation (Miner, Bauerle, & Baldocchi, 2017). *g*_s_ is determined by the stomatal dynamics (opening and closing of stomatal aperture) as well as maximum conductance via epidermal stomata patterning (stomatal density, size, and distribution) (Dow et al., 2014; Lawson & Matthews, 2020; Leakey et al., 2019; Xie et al., 2021). Stomatal patterning more broadly covaries with vein density, leaf width and canopy temperature (Brodribb & Holbrook, 2003; Brodribb & McAdam, 2011; Cano, Sharwood, Cousins, & Ghannoum, 2019; Prakash et al., 2021). But, complexity in the relationships between these traits means that structural-functional relationships controlling *g*_s_ are still not easily predicted. For example, the relationship between *g*_s_ and stomatal density can be positive (Li, Li, Li, & Zhang, 2017; Xu & Zhou, 2008), undetectable (Zhao, Sun, Kjelgren, & Liu, 2015), or negative (Bresta et al., 2018) across different species or pools of intraspecific variation. Likewise, the relationship between *g*_s_ and stomatal complex area can be positive (Galmés et al., 2013; Li et al., 2017) undetectable (Xu & Zhou, 2008; Zhao et al., 2015) or dependent on the shape of the stomatal complex rather than its size (Xie et al. 2021). Therefore, it is of interest to investigate if variation in stomatal patterning contributes to intraspecific variation in the slope parameters of *g*_s_-models.

When vegetation is sampled across biomes at a global scale, *m* or *g*_1_ are lower when water is less available (Lin et al., 2015). This evidence is in line with theoretical expectations (Damour et al., 2010; Lin et al., 2015; Miner and Bauerle 2017), and the widespread observation that plants have greater iWUE when drought-stressed compared to being well-watered (Leakey et al. 2019; Miner et al., 2017). Leaf, vegetation and land surface models often feature a function that lowers *m* or *g*_1_ as a function of plant or soil water status (Anderegg et al., 2017; Klein, 2014; Wolf, Anderegg, & Pacala, 2016). Models have even been developed linking variation in *m* to abscisic acid concentration in the xylem (Gutschick and Simmonneau 2002). However, there are also numerous examples where the slope parameters of *g*s-models have been insensitive to variation in plant or soil water status in the field, unless water stress was extreme (Gimeno et al., 2015; Misson, Panek, & Goldstein, 2004; Xu & Baldocchi, 2003). Miner and Bauerle (2017) reported significantly lower *m* when maize was water stressed in pots. But, the relationships between *m* and plant or soil water status were weak. *m* did not vary over time in field-grown maize, but it is not clear if plant water status changed over the experimental period or not. Structure-function relationships can underpin physiological plasticity of plants across environmental gradients. For example, acclimation to humidity modifies the link between leaf size and the density of veins and stomata (Carins Murphy, Jordan, & Brodribb, 2014). The integration of the BB model of *g*_s_ with a model predicting maximum *g*_s_ from stomatal anatomical traits was a valuable recent advance (Dow, Bergmann, & Berry, 2014). But, studies of variation in stomatal patterning among crop varieties and studies parameterizing *g*_s_ models have generally occurred in isolation of one another. This study aimed to address knowledge gaps about genetic variation in *g*_s_ model parameters across a gradient of water stress by investigating four inbred lines of maize (Zea mays L.) with a range of stomatal densities.

Notably less data describing *g*_s_ model parameters is available from C_4_ crops than other functional groups (Lin et al., 2015; Miner et al., 2017). This is despite the importance of maize, sugarcane, sorghum, switchgrass and other species as sources of food, fuel, fiber and feed (Leakey, 2009). Maize (*Zea mays*) is a model plant for studying the genomics, genetics and physiology of complex traits in C_4_ plants (Buckler et al., 2009). The development of a machine-learning tool to automatically analyze microscopy images of the leaf epidermis facilitated detailed analysis of variation in stomatal patterning across genetically and anatomically diverse maize inbred lines that were consistent over two growing seasons (Xie et al., 2021). Relative to diversity in the species as a whole, B73 and MS71 were identified as inbred lines with moderate (106 mm^−2^) and low stomatal density (88 mm^−2^), respectively (Xie, 2021). When these lines were crossed and self-pollinated, the resulting recombinant inbred lines (RILs) displayed significant transgressive segregation. The RILs with extremely high stomatal density (111 mm^−2^; Z019E0163) and extremely low stomatal density (74 mm^−2^; Z019E0036) were selected and designated RIL2 and RIL1, respectively.

The parameters of the BB and MED models of *g*_s_ were measured for the four genotypes of maize (B73, MS71, RIL1, and RIL2) under five levels of water supply using the *g*_s_-response curve protocol on individual replicate leaves (Wolz et al., 2017). Water availability treatments were generated using an in-field rain-out shelter facility located in the Midwest U.S., which is the world’s primary region of maize production (USDA-FAS, 2020). Variation in *m* and *g*_1_ for different genotypes and water treatments was analyzed in relation to soil water content (SWC), predawn and midday leaf water potential (Ψ_pd_ and Ψ_md_), as well as measures of leaf photosynthetic gas exchange. This allowed the following predictions to be tested: (1) the *m* and *g*_1_ parameters of *g*_s_ models will be lower when water supply is restricted; and (2) the plasticity of *m* and *g*_1_ in response to drought stress will vary among genotypes with distinct stomatal patterning.

## Materials and methods

### Field site and experimental treatments

The study was conducted at a field rain-out shelter facility (Fig. S1) on the University of Illinois at Urbana-Champaign research farm in Champaign, IL, USA (https://www.igb.illinois.edu/soyface/, 40°02’N, 88°14’W) in 2019. The soil type at this site is Drummer–Flanagan series (fine-silty, mixed, mesic Typic Endoa-quoll). It is an organically rich, highly productive Corn Belt soil. The field is tile-drained and has been in continuous cultivation of arable crops for decades. The rain-out shelter had an automatically retractable roof and walls (A-Frame, Cravo Equipment Ltd, Brantford, ON, Canada) that were used to exclude precipitation from a field plot with dimensions of ~76 × 9 m. A weather station integrated with the control system (Igrow 1400, Link4 Corporation, Anaheim, CA) automatically closed the roof and walls within 2 minutes of precipitation being detected by an optical rain sensor. The roof and walls automatically re-opened after no precipitation was detected for 10 minutes. To prevent lateral percolation of water into the soil covered by the rain-out shelter, a plastic barrier impermeable to water was buried vertically from the soil surface to a depth of 1.2 m around the perimeter of the rain-out shelter.

Ten rows of crops were planted along the length of the facility at a spacing of 0.76 m. The outer rows of plants were treated as a border and not sampled, leaving the eight central rows for experimental material. Water was supplied by a surface drip irrigation system (ET256-50SX, Rain Bird Corporation, Azusa, CA) with ten independently controlled zones along the length of the facility. Each irrigation zone was 3 m in length and separated from neighboring zones by alleys of 1.2 m. The zones nearest the end walls of the facility were treated as a border and not sampled. Data was collected for this experiment from plants growing in five zones, which received irrigation once or twice per week over the growing season to achieve totals of 15, 31, 46, 138 and 647 mm (Fig. S2). After sowing, all plots were irrigated to the field capacity to ensure the normal emergence of maize seeds.

Four inbred maize genotypes were studied: two founder lines (B73 and MS71) of the US maize Nested Association Mapping (NAM) population (Buckler et al. 2009) and two recombinant inbred lines (RILs; Z019E0036 and Z019E0163, hereafter referred to as RIL1 and RIL2) resulting from the cross of B73 × MS71 (USDA Germplasm Resources Information Network, GRIN). Within each irrigation zone, each genotype was planted in two rows that were 3 m long with a plant spacing of 16 cm. Planting was performed by hand on June 14th.

### *g*_s_-response curves

The response of steady-state *g*_s_ to variation in photosynthetic photon flux density (PPFD) was measured on the youngest fully expanded leaves near the top of the canopy from August 27 to September 3, 2019, using six portable photosynthetic gas exchange systems (Model Li-6800; Li-Cor Inc., Lincoln, NE, USA) in the field rain-out shelter. The approach of Ball et al. (1987), Leakey et al. (2006), and Wolz et al. (2017) were followed, but on attached leaves in the field. Leaves were acclimated in the chamber with a target leaf temperature of 30.0 °C, relative humidity of 60 %, reference cuvette CO_2_ concentration of 400 μmol mol^−1^, PPFD of 2000 μmol m^−2^ s^−1^, and flow rate of 400 μmol s^−1^ for approximately 30 minutes. Once *g*_s_ had attained steady-state rates, incident PPFD was decreased stepwise to 1500, 1200, 900, 700, 500, 400, 300, 200, 100, and 50 μmol m^−2^ s^−1^. At each light level, *g*_s_ were allowed to reach steady-state before results were recorded (less than 30 minutes) and the next stepwise change was initiated (Fig. S3a). A full *g*_s_-response curve took a minimum of 120 min with 11 points of light levels. Measurements were started around 7 am and continued no later than 2 pm. A total of 71 leaves were measured providing 3-4 replicate samples per genotype in each of the five levels of water supply.

### Leaf water potential, specific leaf area and soil water content

After in situ gas exchange measurements were completed each day, the same leaves were collected to measure leaf water potential (Ψ_md_) using a pressure chamber (PMS Instrument Company; Corvallis, Oregon, USA). Then, four leaf discs (approximately 7.1 cm^2^ per plant) were removed and dried in an oven at 70°C to constant weight and weighed for calculation of specific leaf area (SLA). At predawn the next day, a neighboring leaf from the same plants was collected to measure pre-dawn leaf water potential (Ψ_pd_).

Access tubes were installed within crop rows using a tractor-mounted, customized hydraulic soil corer (Rajurkar et al., 2021) at four locations in each subplot to allow measurement of soil volumetric water content (SWC) twice per week at four depths from 5 to 83 cm with 5-24, 25-43, 44-63, and 64-83 cm, respectively, using the TRIME-PICO TDR system (IMKO GmbH, Ettlingen, Germany).

### Parameterization of stomatal conductance models

Data gathered from the *g*_s_-response curves were used to estimate parameters of the BB (*m*) and MED models (*g*_1_) for each leaf by the least-squares and nonlinear regressions of the following functions (Fig. S3b).

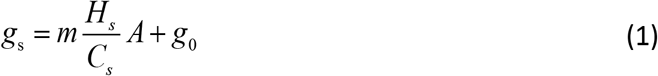

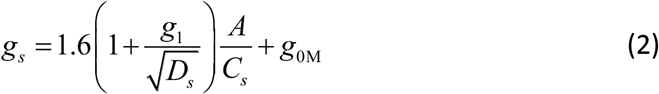

*g*_1_ is defined as:

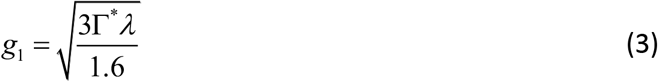

where Γ* is the CO_2_ compensation point for photosynthesis without dark respiration (μmol mol^−1^), and λ the marginal water cost of carbon gain (mmol water μmol^−1^ CO_2_). For similar conditions of temperature and over a moderate range of relative humidity (~40% – 80%), *m* and *g*_1_ are approximately related by the following forms.

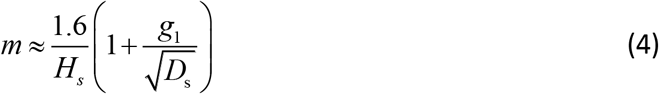

*H*_s_ and *C*_s_ were calculated as:

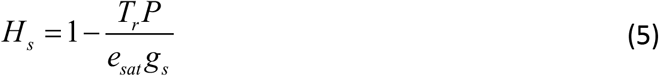

where *T*_r_ is transpiration rate (mol m^−2^ s^−1^), *P* is air pressure (Pa), and *e*_sat_ is the saturated vapor pressure at the substomatal cavity related to leaf temperature (Pa).

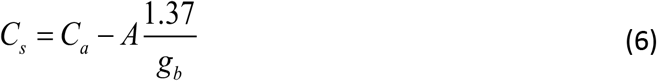

where *C*_a_ is the CO_2_ concentration in the sample chamber (μmol mol^−1^), *g*_b_ is the leaf boundary conductance (mol m^−2^ s^−1^), and 1.37 is the ratio of the molecular diffusivities for H_2_O to CO_2_ at the leaf surface.

The slope parameters *m* and *g*_1_ were obtained by linear and nonlinear fitting to leaf gas exchange data for Equations (1) and (2), respectively. The intercepts, *g*_0_ and *g*_0M_, are often thought to represent the cuticular *g*_s_, or the conductance with closed stomata. Similar to previous studies (Franks et al., 2018; Lin et al., 2015; Wolz et al., 2017; Wu et al., 2020), we did not fit *g*_0_ and *g*_0M_ and taken them as zero due to their uncertainty and correlation with the slopes. Quality assurance was performed by evaluating the goodness-of-fit between the BB model and measured data, with data from all leaves passing the criteria of an R^2^ ≥ 0.9 (Fig S3c).

### Statistical analyses

Differences among four genotypes of maize under a range of water availability were tested by evaluating the slopes and intercepts of the relationships among the BB and MED model parameters (*m* and *g*_1_), leaf water potential (Ψ_md_ and Ψ_pd_), light-saturated gas exchange parameters (*A*_sat_ and *g*_sat_) and mean SWC during the period of physiological data collection, using an analysis of covariance (ANCOVA) test. The null expectation (H0) was that the slope of the regression (or intercept) would not deviate significantly among genotypes. The ANCOVA test was performed using the app “Compare Linear Fit Parameters and Datasets” in OriginPro 2019 (OriginLab Corporation, Northampton, MA, USA). Regression lines were fit for each genotype when either the slopes or intercepts of the relationship between two traits were significantly different among genotypes. Regressions lines were fit across all genotypes when both the slopes and intercepts of the relationship between them were not significantly different among genotypes. To avoid Type II errors, a probability level of *p* = 0.1 was set as the threshold for significance.

## Results

### Responses of *g*_s_-model parameters to varying water availability and plant water status

The slope parameters (*m* or *g*_1_) of *g*_s_-models were lower in value when the average SWC at soil depth profile of 5-83 cm was lower (Fig. 1a,b, *p* < 0.002; Table S1). But, there were no significant differences among genotypes in the relationship between SWC and *m* or *g*_1_ (Fig. 1a,b; *p* > 0.662). This consistency in *m* and *g*_1_ among genotypes as SWC varied with irrigation rate was observed regardless of the soil depth at which SWC was measured (Fig. 1c-j). However, the proportion of variance in *m* or *g*_1_ explained by SWC varied from being strongest for intermediate soil depths (44-63 cm; R^2^ = 0.50-0.52; Fig. 1g,h) followed by deeper soil layers (64-83 cm; R^2^ = 0.43-0.44; Fig. 1i,j) and shallower soil layers (5-24 cm and 25-43 cm; R^2^ = 0.36-0.37; Fig. 1c-f). And, this corresponded with *m* or *g*1 being more sensitive to a given change in SWC at intermediate and deeper soil depths than the equivalent changes in SWC in shallower soil layers (Fig. 1).

**Fig. 1.**
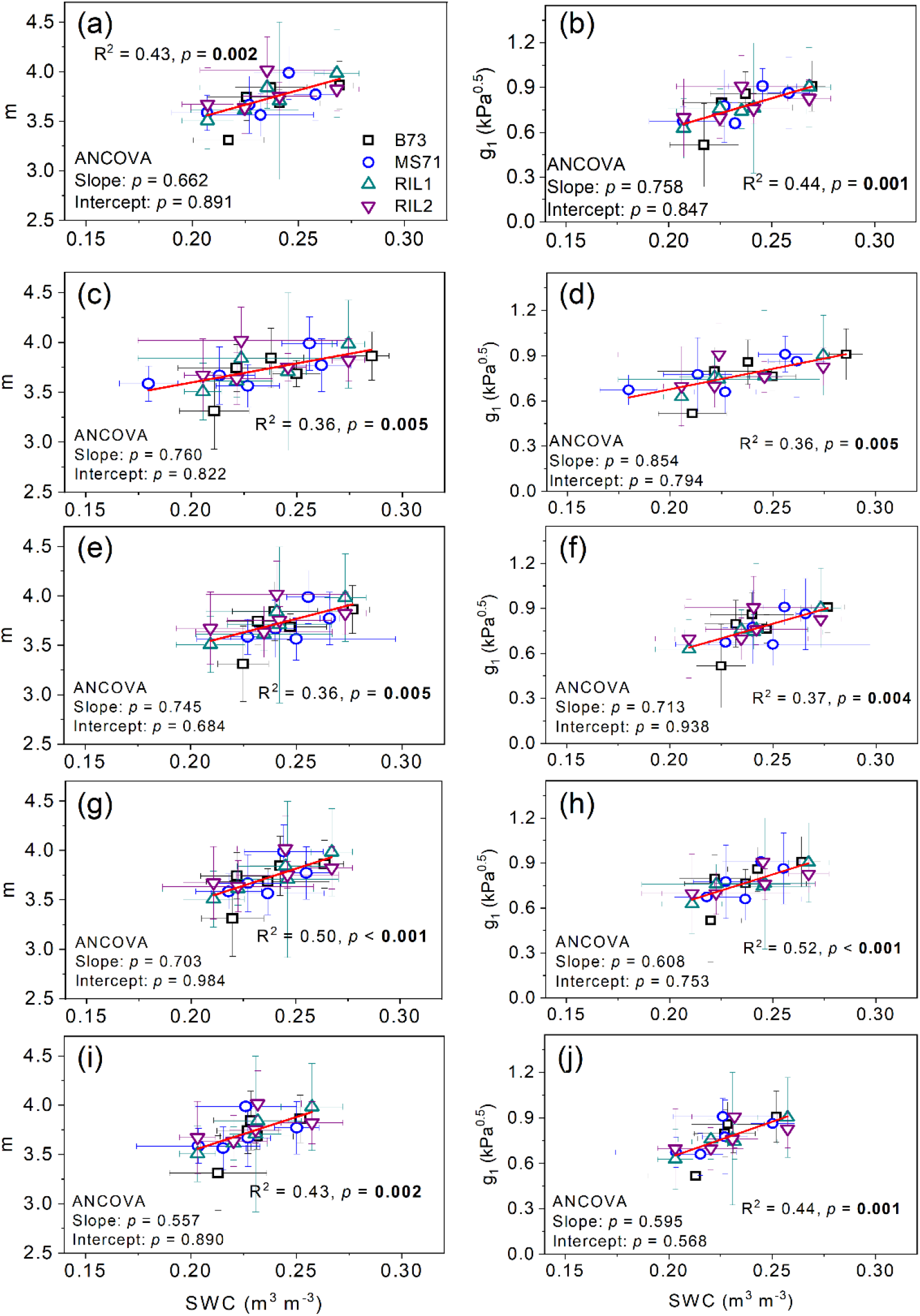
Relationships between the BB slope (*m*) and MED slope (*g*_1_) with average soil water content (SWC) at depths of 5-83 cm (a,b), 5-24 (c,d), 25-43 (e,f), 44-63 (j,h), and 64-83 cm (i,j) for four genotypes of maize (B73, MS71, RIL1, and RIL2) under five levels of water supply. The statistical significance of genotypic variation in the slope or intercept of the response to SWC (ANCOVA) is inset, along with the results of correlation analysis for the group of genotypes or single genotypes, as appropriate. Plotted points are genotype means at each level of SWC ± SD.

Plant water status across the range of SWC was characterized in terms of leaf water potential both pre-dawn (Ψ_pd_) and during the mid-day period (Ψ_md_). *m* and *g*_1_ both were lower when Ψ_pd_ was more negative (Fig. 2a,b). But, the relationships of *m* or *g*_1_ with Ψ_md_ were only marginally significant (Fig. 2c,d) and neither Ψ_pd_ or Ψ_md_ explained as much variation in *g*_s_-model slope parameters as SWC. There was also no variation among genotypes in these relationships between *m* or *g*_1_ and Ψ_pd_ or Ψ_md_ (Fig. 2; *p* > 0.103).

**Fig. 2.**
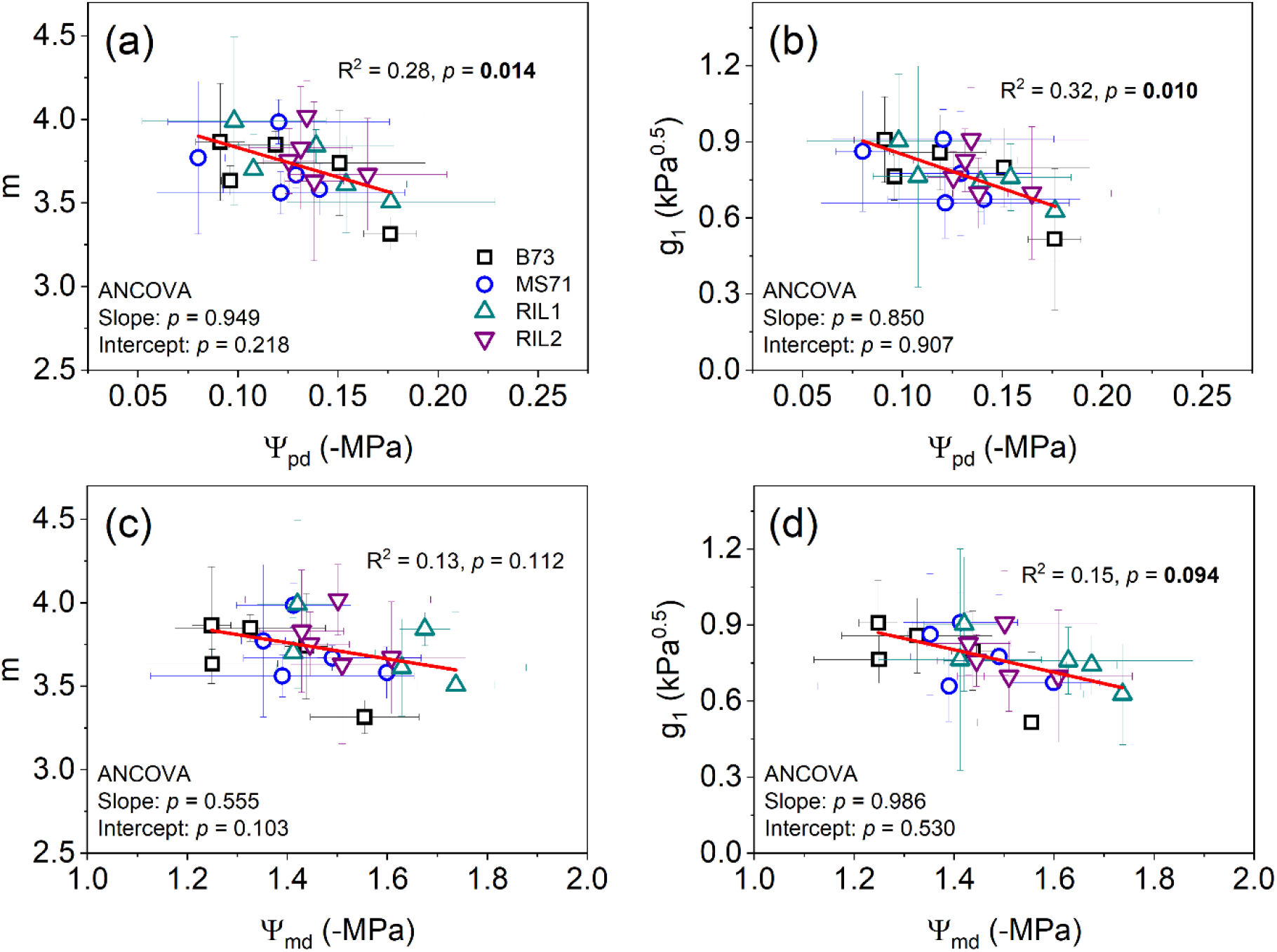
The relationships between the BB slope (*m*) and MED slope (*g*_1_) with predawn and midday leaf water potential (Ψ_pd_ and Ψ_md_) for four genotypes of maize (B73, MS71, RIL1, and RIL2) under five levels of water supply. The results of statistical tests are provided as described in Fig. 1. Plotted points are genotype means at each level of SWC ± SD.

### *A*_sat_, *g*_sat_ and SLA as drivers of variation in *g*s-model parameters under varying SWC

Drought-induced variation in *m* was significantly associated with variation in both *g*_sat_ and *A*_sat_ in a genotype-specific fashion (Fig. 3a,b, *p* < 0.049). The sensitivity of *m* or *g*_1_ to *g*_sat_ or *A*_sat_ were consistent across all genotypes (i.e., regression slopes did not significantly differ, *p* > 0.445), but the value of *m* of *g*_1_ for a given *g*_sat_ or *A*_sat_ differed between genotypes (i.e., regression intercepts significantly varied, *p* ≤ 0.049). The proportion of variation in *g*_s_-model parameters explained by *g*_sat_ and *A*_sat_ was very similar, as described by goodness-of-fit, i.e., R^2^ (Fig. 3). These relationships stem from lower SWC driving progressively lower *g*_sat_ and *A*_sat_ in a genotype-specific manner (*p* < 0.013, Fig. 4a,b). And, the observed intercept changes in *g*_sat_ or *A*_sat_ were driven by the genotype-specific responses to SWC. Drought-induced variation in *g*_sat_ and *A*_sat_ was significantly associated with Ψ_md_ (R^2^ > 0.67, *p* < 0.001; Fig. 4c,d), i.e., *g*_sat_ and *A*_sat_ covaried with leaf water status assessed immediately after gas exchange measurements were completed. And, this response was consistent across all four genotypes (*p* > 0.201). The correlations between *g*_s_ and Ψ_pd_ as well as *A* and Ψ_pd_ were species-specific (*p* < 0.051), with genotypes having different *g*_s_ of *A* during the day even when the water status of the plants had been equivalent pre-dawn (Fig. 4e,f). Neither *m* nor *g*_1_ was significantly correlated with SLA (Fig. 5a,b, *p* > 0.604).

**Fig. 3.**
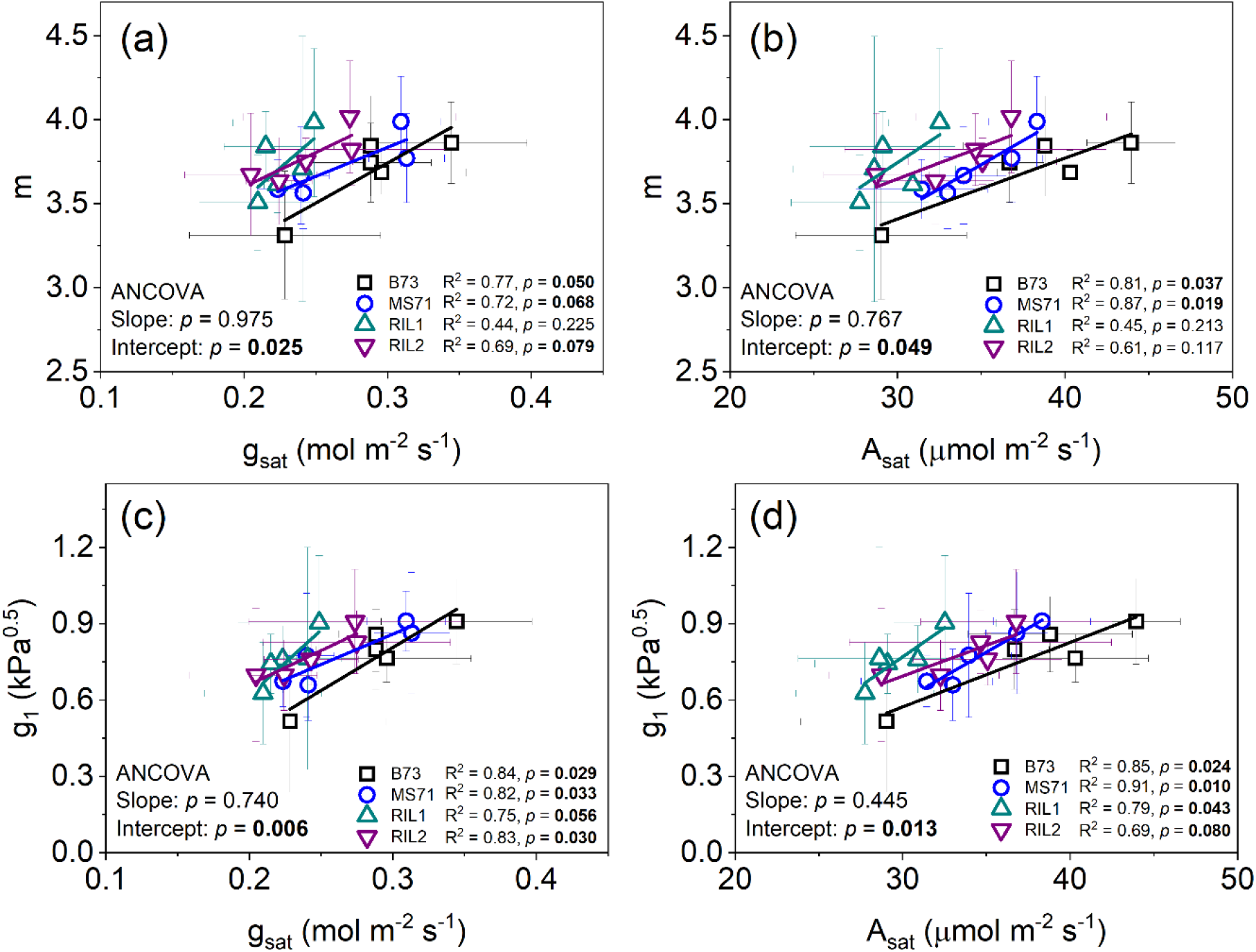
The relationships between the BB slope (*m*) and MED slope (*g*_1_) with light-saturated photosynthesis rate (*A*_sat_) and stomatal conductance (*g*_sat_) for four genotypes of maize (B73, MS71, RIL1, and RIL2) under five levels of water supply. The results of statistical tests are provided as described in Fig. 1. Plotted points are genotype means at each level of SWC ± SD.

**Fig. 4.**
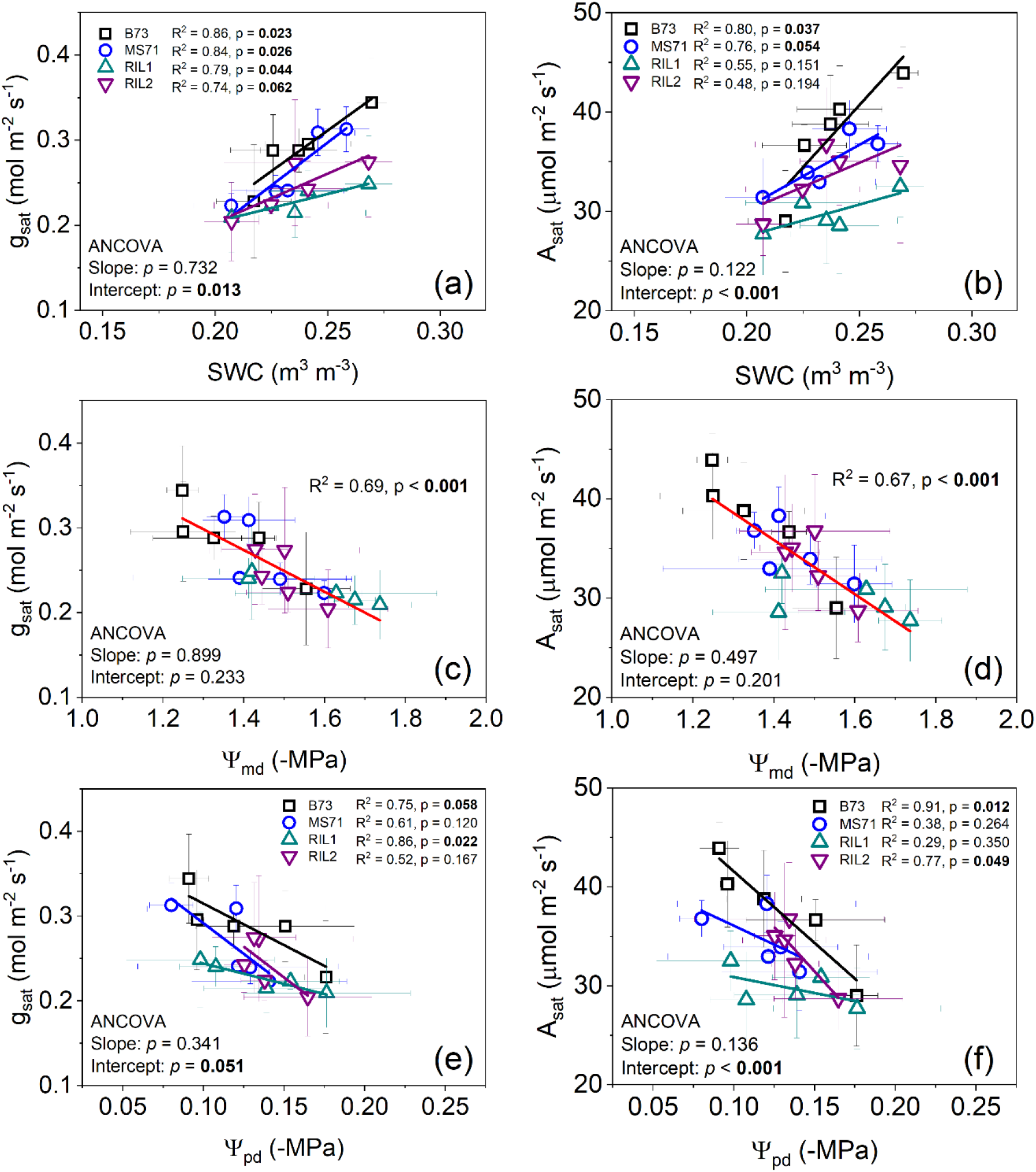
The relationships between light-saturated photosynthesis rate (*A*_sat_) and stomatal conductance (*g*_sat_) with average soil water content (SWC) at a soil depth of 5-83 cm (a,b), and predawn and midday leaf water potential (Ψ_pd_ and Ψ_md_, c-f) for four genotypes of maize (B73, MS71, RIL1, and RIL2) under five levels of water supply. The results of statistical tests are provided as described in Fig. 1. Plotted points are genotype means at each level of SWC ± SD.

**Fig. 5.**
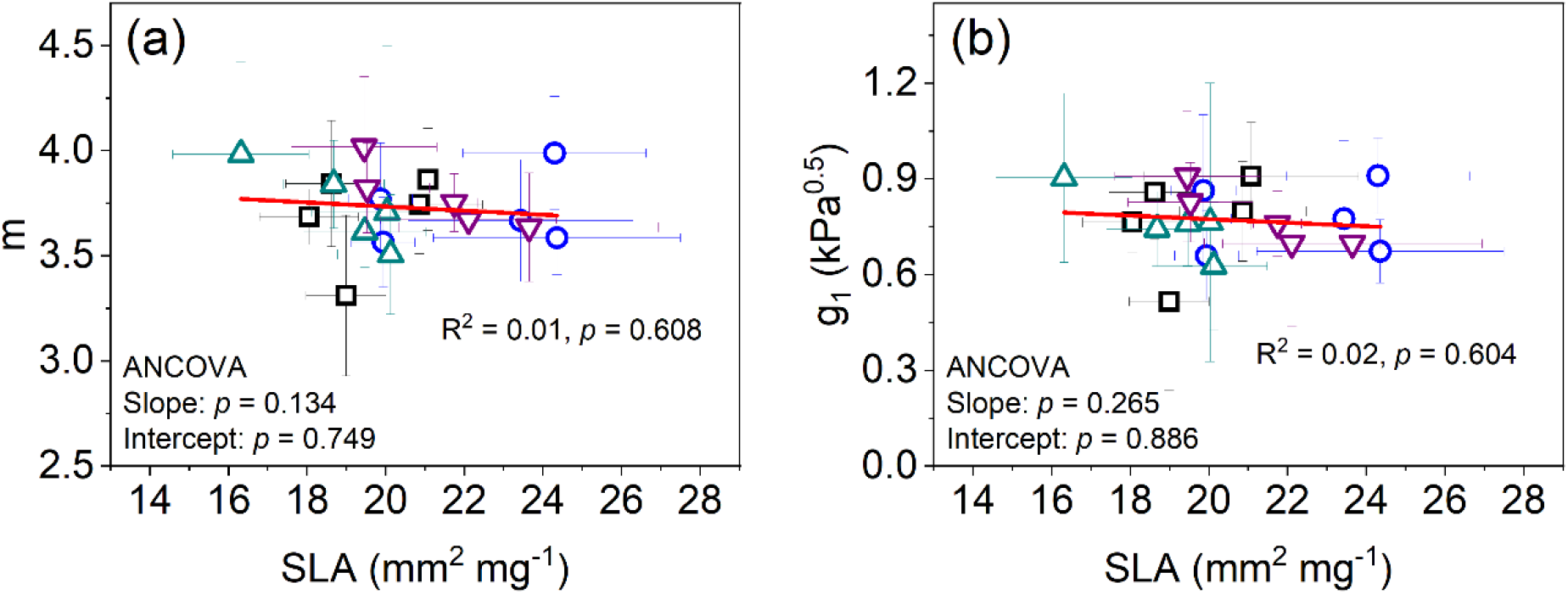
The relationships between the BB slope (*m*) and MED slope (*g*_1_) with specific leaf area (SLA) for four genotypes of maize (B73, MS71, RIL1, and RIL2) under five levels of water supply. The results of statistical tests are provided as described in Fig. 1. Plotted points are genotype means at each level of SWC ± SD.

## Discussion

This study successfully addressed its aims to investigate how genotypic variation in *g*_s_-model parameters among four anatomically distinct maize inbred lines were impacted by a gradient in water availability. As predicted, *m* and *g*_1_ were progressively lower when plants were more drought stressed due to withholding of water supply (Fig. 1). Variation in *m* and *g*_1_ showed the strongest relationships with water availability in deeper soil layers, moderate dependency on Ψ_pd_, and only a marginally significant association with Ψ_md_ (Figs. 1,2). Contrary to expectations, inbred genotypes of maize that significantly vary in stomatal patterning, *g*_s_ and A (Figs. 3,4) were very consistent with respect to *g*_s_-model parameters and their plasticity in response to drought stress (Fig. 1). These findings provide new evidence to guide how models of maize should simulate *g*_s_ and its influence on plant function across a range of water status that is relevant to field conditions in the primary growing region of this major crop.

The line of best fit describing how *m* varies with SWC across the whole rooting zone (Fig. 1a) was very similar to that for pot-grown maize (Miner and Bauerle 2017). But, this probably is somewhat coincidental because the physical characteristics of the soils in the two experiments are very different, so the moisture release curves and effects on plant water status of the two gradients in water supply were expected to differ. This interpretation is consistent with the well-watered greenhouse grown plants having Ψ_md_ equivalent to Ψ_pd_ in the field, but much less negative than Ψ_md_ in the field (Fig. 2; Miner and Bauerle 2017). Nonetheless, the consistency in the direction of response in the two studies, and the consistency in response among anatomically diverse inbred lines in the present study, suggests that the results do provide a reasonable first approximation of how maize stomata operate in a production setting and how that should be parameterized in models.

The data presented here are valuable because there are far fewer estimates of slope parameters for *g*_s_ models (i.e., *m* and *g*_1_) for C_4_ species than C_3_ species in general, and especially under field conditions (Lin et al. 2015; Miner et al., 2017). On average, *m* was 3.87 and *g*_1_ was 0.88 kPa^0.5^ across the four genotypes of maize under well-watered conditions (Fig. 1, Table S1). This sits between parameter estimates previously published for maize grown in controlled environment conditions (*m* = 3.06, Ball 1988; *m* = 3.23, Collatz et al., 1992; *m* = 4.53, Miner and Bauerle 2017; *g*_1_ = 1.281; Yun *et al.*, 2020) and very close to a parameter estimate for maize in the field in Colorado (*m* = 3.72, Miner and Bauerle 2017). The results are also comparable to measurements of *Panicum virgatum* (*m* = 3.9), *Miscanthus x giganteus* (*m* = 3.3) and *Sorghum bicolor* (*m* = 4.32) grown at nearby field sites (LeBauer et al., 2013; Li et al., 2021) as well as C_4_ grasses in general (*m* = 4.1, Miner *et al.*, 2017; *m* = 4.0, Franks et al., 2017). But, relatively subtle variation in *g*_*s*_-model parameters can significantly impact predictions of leaf, canopy, ecosystem and global water fluxes (Franks et al., 2017; Wolz et al., 2017), so additional data collection is still needed. Investigation of hybrid maize as well as maize lines that capture additional genetic and physiological diversity would be particularly valuable to aid in simulations of carbon and water fluxes for this key crop and the U.S. Corn Belt region as a whole.

There is significant uncertainty surrounding the physiological mechanisms that underpin variation in *m* or *g*_1_ across different growing conditions in either time or space (Damour et al., 2010; Héroult, Lin, Bourne, Medlyn, & Ellsworth, 2013; Miner at al., 2017; Xu & Baldocchi, 2003). Some studies demonstrated that *m* and *g*_1_ are relatively stable under drought conditions (Gimeno et al., 2016) or the inclusion of leaf water potential did not improve model performance (Wu et al., 2020). In a natural oak-grass savanna, *m* for blue oak remained constant through a severe summer drought (Xu & Baldocchi, 2003). But, others have found a response in *g*_s_-model parameters to water-deficit (Anderegg et al., 2017; Damour et al., 2010; Sellers et al., 1996; Venturas et al., 2018). In a common garden experiment, *m* decreased under drought in two *Eucalyptus* species from humid regions but not in two other eucalypts from drier regions (Héroult et al., 2013). This mechanistic uncertainty is reflected in a subset of models variously using SWC, soil Ψ, plant water status, or even hormone concentrations to modulate simulations of stomatal behavior in response to drought stress (Anderegg et al., 2017; Damour et al., 2010; Oleson et al., 2010; Sellers et al., 1996; Sperry et al., 2017; Venturas et al., 2018). Variations in *m* or *g*_1_ for field-grown maize most closely correlated with SWC in intermediate to deep layers of the rooting profile, were moderately correlated with Ψ_pd_, and most weakly correlated with Ψ_md_ (Figs. 1,2). Ψ_pd_ is commonly considered to be in equilibrium with soil Ψ and the observed data indicate that the water supply treatments here caused long-term variation in soil water status that was beyond the capacity of the system to recover overnight. Nevertheless, the differences between Ψ_pd_ and Ψ_md_ indicate that significant additional short-term water stress did develop during the day as the evaporative demand of the crop was met to differing degrees at the different levels of water supply. This strong role of water stress that temporarily develops during the day is evident from the relationships of *g*_s_ and *A* with Ψ_md_ than Ψ_pd_ (Fig. 4). But, crucially, the weaker relationship of *m* or *g*_1_ with Ψ_md_ than Ψ_pd_ implies that the plasticity in parameters of *g*_s_ models is driven by long-term signals and responses rather than the short-term responses to drought within a single day. This may include changes in photosynthetic capacity, which can influence the sensitivity of *g*_s_ to atmospheric conditions (Franks et al., 2017), but further work will be needed to resolve the mechanistic details. Notably, no relationship was found between SLA and *m* or *g*_1_ among the four maize inbred lines (Fig. 5). This contrasts with the results of a study of tropical rainforest trees, but may reflect the consequence of studying intraspecific rather than interspecific variation in traits (Wu et al. 2020).

The four maize inbred lines studied display significant variation in stomatal density, other aspects of stomatal patterning and anatomy (Xie et al. 2021), and *A* and *g*_s_ (Fig. 4). Nevertheless, they had very similar *m* and *g*_1_ (Figs. 1,2). And, genotype-specific plasticity in *m* or *g*_1_ in response to a gradient of water supply could not be detected. This convergence in *g*_s_-model parameter values across genotypes is consistent with the trade-off between carbon gain and water use (i.e., WUE) being the most fundamental trade-off for terrestrial plant life (Briggs and Shantz 1917; Boyer et al., 1982; Hetherington and Woodward 2003). And, it indicates that there must be significant flexibility in structure-function relationships between stomatal patterning and other aspects of leaf gas exchange. Or in other words, the same WUE can be achieved with different configurations of stomata, leaf hydraulics and photosynthesis. This complements emerging frameworks for understanding the tight coordination in the photosynthetic, gas exchange and water supply capacities of leaves across the diversity of land plants (Deans, Brodribb, Busch, & Farquhar, 2020). It is also important to recognize that flexibility in structure-function relationships of the type observed here will set constraints and maybe create opportunities for efforts to engineer or select for improved crop WUE (Leakey et al., 2019). New high-throughput phenotyping and analytical techniques are providing unprecedented detail and depth of information about the suite of traits that underpin variation in WUE within C_4_ species (Ferguson et al., 2021; Pignon et al., 2021a,b; Xie et al., 2021). This should then in turn allow additional studies of the type presented here to quantify *g*_s_-model parameters in other genotypes and provide the parameterization data needed to inform crop improvement effort with *in silico* analyses (Marshall-Colon et al., 2017).

## Acknowledgements

We thank Luke Freyfogle and Yu Wang for technical and field assistance, and Shuai Li for assistance with data analysis.

