## Supplemental Figs for "Plasticity in stomatal behavior across a gradient of water supply is consistent among field-grown maize inbred lines with varying stomatal patterning"

**SUPPORTING INFORMATION**

**Table S1. Leaf-level slope estimates of the BB model (*m*) and MED model (*g*_1_) and goodness-of-fit (R^2^) for four genotypes of maize (B73, MS71, RIL1, and RIL2)** **under five levels of water supply (W).**

| **Water supply** | **Genotypes** | **Replicates** | ***m*** | **R^2^** | ***g*_1_** |
| --- | --- | --- | --- | --- | --- |
| W1 | B73 | 1 | 3.72 | 0.990 | 0.79 |
| W1 | B73 | 2 | 3.55 | 0.969 | 0.71 |
| W1 | B73 | 3 | 2.71 | 0.995 | 0.07 |
| W1 | B73 | 4 | 3.27 | 0.993 | 0.49 |
| W1 | MS71 | 1 | 3.38 | 0.989 | 0.55 |
| W1 | MS71 | 2 | 3.57 | 0.993 | 0.68 |
| W1 | MS71 | 3 | 3.81 | 0.966 | 0.79 |
| W1 | RIL1 | 1 | 3.91 | 0.947 | 0.93 |
| W1 | RIL1 | 2 | 3.51 | 0.989 | 0.63 |
| W1 | RIL1 | 3 | 3.11 | 0.986 | 0.37 |
| W1 | RIL1 | 4 | 3.49 | 0.970 | 0.58 |
| W1 | RIL2 | 1 | 4.07 | 0.954 | 1.03 |
| W1 | RIL2 | 2 | 4.00 | 0.989 | 0.88 |
| W1 | RIL2 | 3 | 3.33 | 0.988 | 0.47 |
| W1 | RIL2 | 4 | 3.29 | 0.985 | 0.42 |
| W2 | B73 | 1 | 3.41 | 0.997 | 0.58 |
| W2 | B73 | 2 | 3.77 | 0.986 | 0.81 |
| W2 | B73 | 3 | 3.74 | 0.987 | 0.79 |
| W2 | B73 | 4 | 4.07 | 0.979 | 1.02 |
| W2 | MS71 | 1 | 4.07 | 0.997 | 1.12 |
| W2 | MS71 | 2 | 3.54 | 0.996 | 0.65 |
| W2 | MS71 | 3 | 3.39 | 0.994 | 0.56 |
| W2 | RIL1 | 1 | 3.61 | 0.989 | 0.86 |
| W2 | RIL1 | 2 | 3.82 | 0.994 | 0.84 |
| W2 | RIL1 | 3 | 3.41 | 0.987 | 0.57 |
| W2 | RIL2 | 1 | 4.05 | 0.994 | 0.91 |
| W2 | RIL2 | 2 | 3.42 | 0.991 | 0.59 |
| W2 | RIL2 | 3 | 3.65 | 0.986 | 0.73 |
| W2 | RIL2 | 4 | 3.42 | 0.968 | 0.56 |
| W3 | B73 | 1 | 3.76 | 0.983 | 0.85 |
| W3 | B73 | 2 | 4.25 | 0.996 | 1.04 |
| W3 | B73 | 3 | 3.53 | 0.987 | 0.68 |
| W3 | MS71 | 1 | 3.60 | 0.993 | 0.70 |
| W3 | MS71 | 2 | 3.29 | 0.993 | 0.47 |
| W3 | MS71 | 3 | 3.81 | 0.984 | 0.81 |
| W3 | RIL1 | 1 | 3.90 | 0.977 | 0.78 |
| W3 | RIL1 | 2 | 3.51 | 0.983 | 0.55 |
| W3 | RIL1 | 3 | 4.09 | 0.976 | 0.85 |
| W3 | RIL1 | 4 | 3.87 | 0.986 | 0.80 |
| W3 | RIL2 | 1 | 4.18 | 0.977 | 1.04 |
| W3 | RIL2 | 2 | 3.61 | 0.991 | 0.63 |
| W3 | RIL2 | 3 | 3.81 | 0.989 | 0.80 |
| W3 | RIL2 | 4 | 4.48 | 0.967 | 1.16 |
| W4 | B73 | 1 | 3.84 | 0.965 | 0.89 |
| W4 | B73 | 2 | 3.52 | 0.992 | 0.66 |
| W4 | B73 | 3 | 3.70 | 0.993 | 0.75 |
| W4 | MS71 | 1 | 3.64 | 0.994 | 0.76 |
| W4 | MS71 | 2 | 4.04 | 0.980 | 0.94 |
| W4 | MS71 | 3 | 4.29 | 0.951 | 1.04 |
| W4 | RIL1 | 1 | 3.55 | 0.911 | 0.77 |
| W4 | RIL1 | 2 | 4.75 | 0.962 | 1.29 |
| W4 | RIL1 | 3 | 2.83 | 0.991 | 0.23 |
| W4 | RIL2 | 1 | 3.76 | 0.977 | 0.80 |
| W4 | RIL2 | 2 | 3.62 | 0.989 | 0.62 |
| W4 | RIL2 | 3 | 3.98 | 0.975 | 0.90 |
| W4 | RIL2 | 4 | 3.66 | 0.986 | 0.72 |
| W5 | B73 | 1 | 3.67 | 0.987 | 0.80 |
| W5 | B73 | 2 | 4.23 | 0.986 | 1.19 |
| W5 | B73 | 3 | 3.92 | 0.991 | 0.90 |
| W5 | B73 | 4 | 3.63 | 0.985 | 0.75 |
| W5 | MS71 | 1 | 4.02 | 0.968 | 1.14 |
| W5 | MS71 | 2 | 3.40 | 0.978 | 0.56 |
| W5 | MS71 | 3 | 3.89 | 0.986 | 0.89 |
| W5 | RIL1 | 1 | 3.49 | 0.980 | 0.63 |
| W5 | RIL1 | 2 | 4.70 | 0.983 | 1.34 |
| W5 | RIL1 | 3 | 3.84 | 0.981 | 0.83 |
| W5 | RIL1 | 4 | 3.90 | 0.972 | 0.83 |
| W5 | RIL2 | 1 | 3.74 | 0.969 | 0.84 |
| W5 | RIL2 | 2 | 3.57 | 0.959 | 0.63 |
| W5 | RIL2 | 3 | 3.84 | 0.983 | 0.86 |
| W5 | RIL2 | 4 | 4.15 | 0.903 | 0.97 |

Fig. S1. Overview of the experimental site and plot layout. (a) a panoramic view of field rain-out shelter facility in a field plot with dimensions of ~76 x 9 m and (b) top view of partial plot layout.

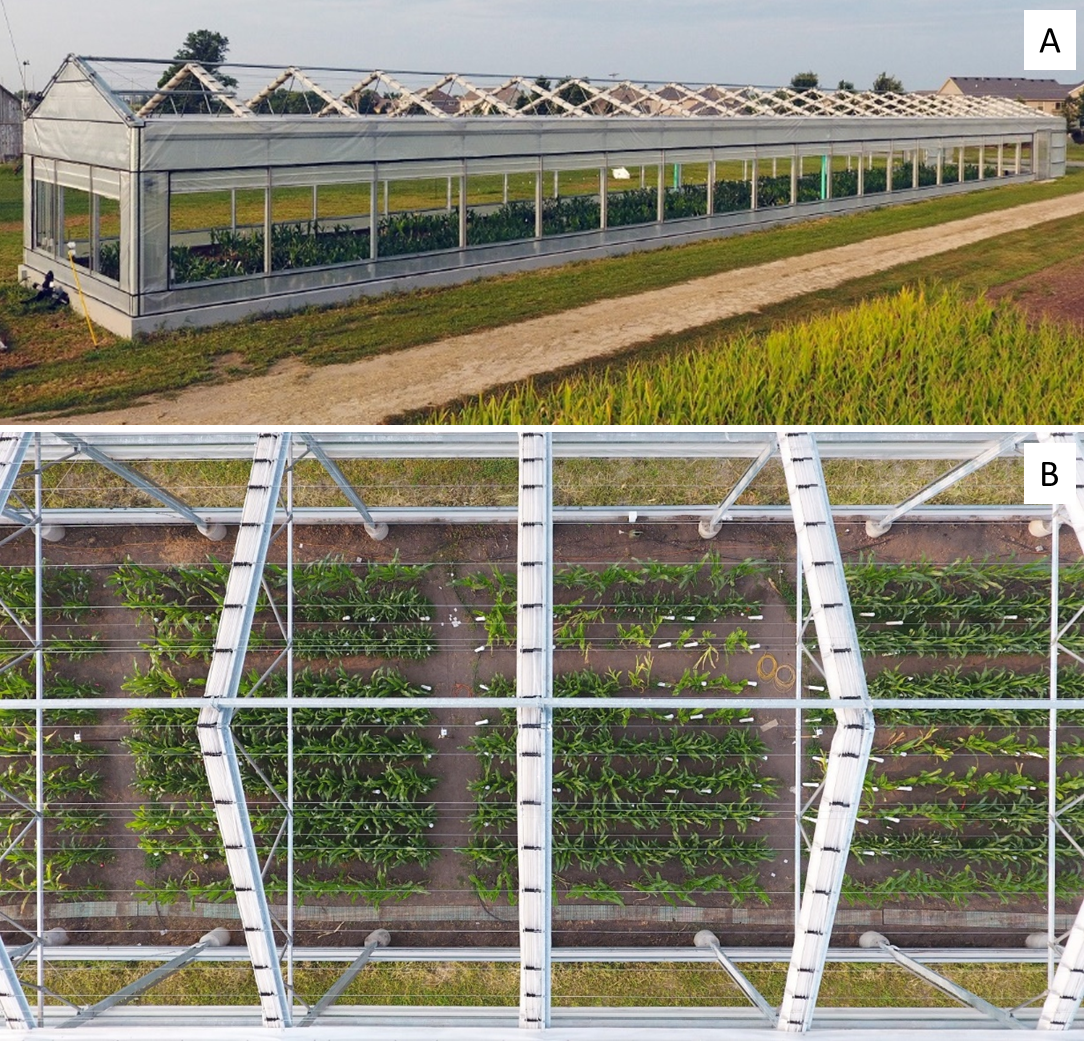

Fig. S2. Total irrigation throughout the whole growing season and rainfall from June to September under five levels of water supply (W).

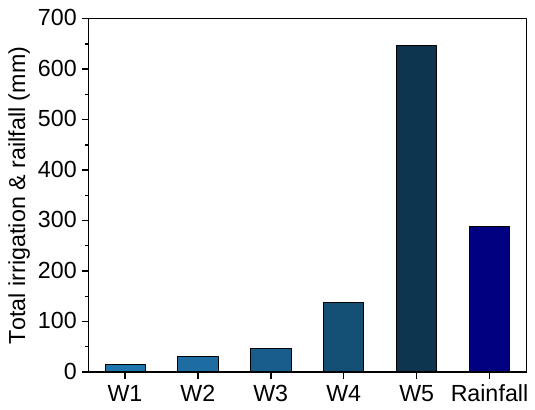

Fig. S3. (a) stable time of stomatal conductance (*g*_s_) for different light levels for the reference maize line (B73) under well-watered regime; (b) relationship of *g*_s_ with the BB Index (*AH*_s_/*C*_s_) for youngest fully expanded leaves of well-watered maize for four genotypes (B73, MS71, RIL1 and RIL2). Regression lines are fit across all data points for each genotypic leaf with three or four replicates; and (c) frequency distribution of the goodness-of-fit (R^2^) when estimating the parameters *m* in the BB model of *g*_s_ from data collected as *g*_s_ -response curves from individual leaves (n = 71) of four genotypes of maize for five water treatments.

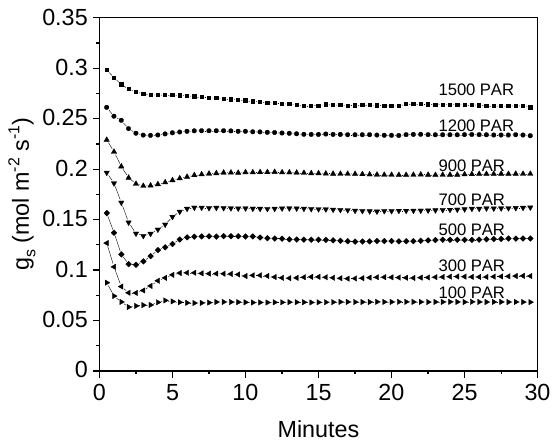

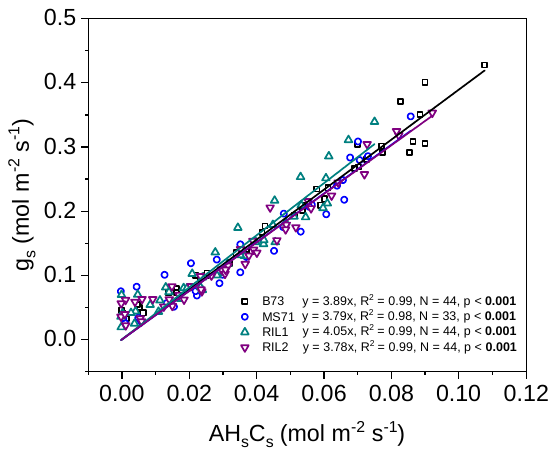

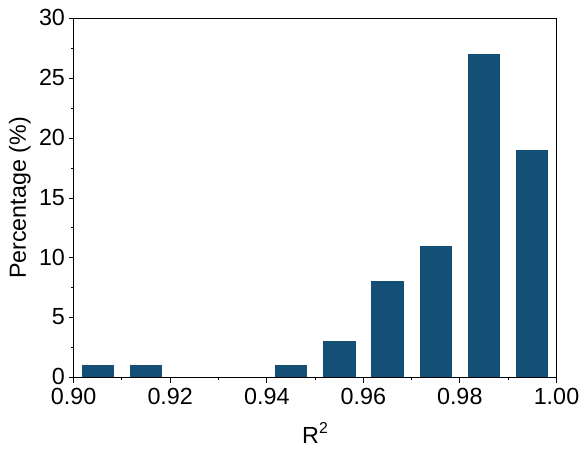

(b)

(a)

(c)
